# Transcriptional clusters follow a conserved condensation-dispersal sequence during stem cell differentiation

**DOI:** 10.1101/2023.07.04.547621

**Authors:** Tim Klingberg, Irina Wachter, Agnieszka Pancholi, Yomna Gohar, Priya Kumar, Marcel Sobucki, Elisa Kämmer, Süheyla Eroğlu-Kayıkçı, Sylvia Erhardt, Carmelo Ferrai, Vasily Zaburdaev, Lennart Hilbert

**Affiliations:** Department of Biology, Friedrich-Alexander-Universität Erlangen-Nürnberg, 91058 Erlangen, Germany; Max-Planck-Zentrum für Physik und Medizin, 91058 Erlangen, Germany; Zoological Institute, Karlsruhe Institute of Technology, 76131 Karlsruhe, Germany; Institute of Biological and Chemical Systems – Biological Information Processing, Karlsruhe Institute of Technology, 76344 Eggenstein-Leopoldshafen, Germany; Institute of Pathology, University Medical Center Göttingen, 37075 Göttingen, Germany

**Keywords:** Nuclear organization, biomolecular condensates, gene expression, enhancer-promoter interaction

## Abstract

Spatiotemporal organization of transcription is essential for organism development. Most eukaryotic genes are transcribed by RNA polymerase II (Pol II). In stem cells, Pol II forms prominent clusters, which gradually disappear during differentiation, such that only smaller clusters remain. Here, we ask whether the formation and loss of large Pol II clusters is a stereotypical process explicable by changes in the Pol II transcriptional state during differentiation. We assess clusters by super-resolution microscopy in differentiating mouse embryonic stem cells, sperm precursor formation in fruit flies, and germ layer induction in zebrafish. In all cases, Pol II clusters first become larger and rounder, then unfold, and finally disperse into small clusters. These shape changes are accompanied by initial increase in recruited Pol II, subsequent transition into transcript elongation, and finally reduction of active enhancers. We reproduce these observations using a biophysical surface condensation model, where enhancers support Pol II cluster formation, and transcriptional activity unfolds clusters. Our work indicates that changes in enhancer marks and transcriptional activity during differentiation define a stereotyped trajectory through a generally applicable space of cluster shapes.

**Graphical Abstract:** 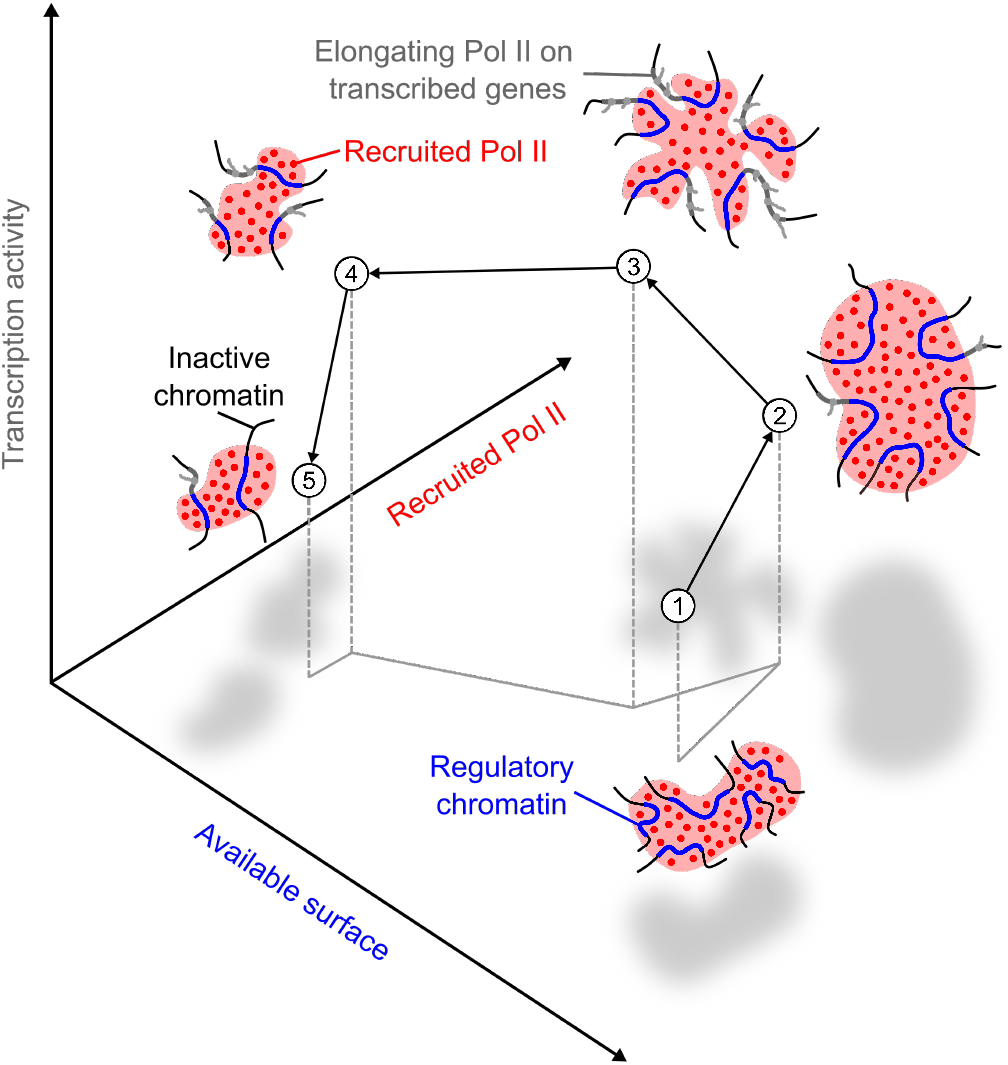

## Introduction

Spatial and temporal control over the transcription of genes into RNA is fundamental to ensure correct development. Altered transcriptional control may lead to developmental defects as well as the emergence of disease in the adult organism. The key regulatory steps of the transcription of genes by the eukaryotic RNA polymerase II (Pol II) are localized within distinct, but adjacently placed spatial compartments on the scale of approximately 100 nm [1, 2, 3, 4]. During the initial recruitment to gene promoters, Pol II forms dynamic macromolecular clusters [5, 6, 7]. As Pol II transitions from the recruited state towards actual elongation of RNA transcripts, a translocation from recruitment-associated transcriptional clusters to distinct compartments associated with RNA splicing [8, 9, 10], or even the dispersal of clusters [7, 11] are observed. Transcriptional clusters, sometimes also referred to as transcription factories [12, 13, 14, 15, 16], have been observed in human cells [17, 12, 13, 18, 6, 19, 20, 21, 16], but also in models of mammalian pluripotency, such as mouse embryonic stem cells (mESCs) [22, 23], as well as models of embryonic development, such as Drosophila (fruit fly) [24, 25, 26] or zebrafish [27, 28, 29, 30, 31, 32]. In differentiated cells, transcriptional clusters persist only for approximately 10 s and contain only several tens of Pol II complexes, whereas in pluripotent cells long-lived, prominent clusters containing hundreds of Pol II complexes can be seen [6, 7, 23, 30]. These long-lived clusters contain locally elevated concentrations of Pol II in the recruited state, whereas elongating Pol II is excluded, resulting in a variety of cluster shapes [30]. High levels of Pol II recruitment and tightly controlled transitions from a recruited to an elongating state have also been implicated in embryonic development and differentiation of stem cells [33, 34, 35, 36, 37].

Mechanistically, the clustering of Pol II and transcription factors associated with the different steps of transcriptional control has been attributed to liquid-liquid phase separation processes [38, 23, 39, 8, 40, 41, 42, 43]. In the case of the more prominent clusters seen in stem cells, a more specific scenario has been proposed, where regulatory genomic regions act as binding surfaces that locally facilitate the formation of liquid condensates [30]. More generally speaking, surface condensation refers to a mechanism that allows formation of condensates below the saturation concentration required for canonical liquid-liquid phase separation, exploiting, for example, DNA or chromatin as a microscopic surface that locally increases concentrations of the phase-separating species [44, 45]. In pluripotent embryonic cells, regulatory chromatin regions such as super-enhancers were proposed to serve as condensation surfaces underlying the formation of clusters enriched in recruited Pol II [30]. This theory is in line with findings that super-enhancers are frequently associated with cellular pluripotency and are central to transcriptional control in development [46, 47, 48, 49].

In mESCs that undergo differentiation, the loss of prominent transcriptional clusters occurs gradually [23], indicating a continuous transition from a stem-cell-specific organization of Pol II towards an organization that is more characteristic of a differentiated cell. In this study, we ask in how far changes in the organization of Pol II clusters over the course of differentiation can be explained in terms of surface condensation at super-enhancers. Assessing three different models of stem cell differentiation (pluripotency exit of mouse embryonic stem cells, fruit fly sperm precursor formation, and germ layer induction in zebrafish embryos), we find that the regulatory state of transcriptional clusters follows a stereotypic trajectory with progressing differentiation. In particular, a transient increase in Pol II recruitment is followed by a transient increase in elongation and, finally, a partial loss of the active-enhancer-associated chromatin mark histone 3 lysine 27 acetylation (H3K27ac). We observed a progression characterized by transient growth, unfolding, and subsequent dispersal of prominent Pol II clusters, which was conserved in the three models of stem cell differentiation. This formation-dispersal sequence was equally reproduced by a theoretical coarse-grained surface condensation model, which accepts the levels of recruited Pol II, elongating Pol II, and available super-enhancer surfaces as an input and produces Pol II cluster shapes as an output. These results imply that surface condensation at regulatory chromatin can coherently explain a formation-dispersal sequence of prominent transcriptional clusters that occurs during major transitions in cell identity, seen across different types of stem cell differentiation processes in three different species. Our work therefore provides a biophysical understanding of how differentiating cells assemble and subsequently dismantle transcriptional clusters, elucidating a central process in the control of developmental transcriptional programs.

## Results

### Prominent transcriptional clusters form and disperse in a step-wise transition upon differentiation of cultured mouse embryonic stem cells

We aimed to characterize the formation and loss of transcriptional clusters in different scenarios of stem cell differentiation. To monitor the shape as well as the state of Pol II during the differentiation process, we used differential serine phosphorylation patterns of the C-terminal heptad repeat domain (CTD, amino acid sequence YSPTSPS) of the RNA polymerase II subunit 1. In particular, the recruitment of the inactive unphosphorylated Pol II under involvement of the protein Mediator is followed by phosphorylation of the Serine 5 (Ser5P) of the CTD by cyclin-dependent kinase 7 (CDK7), whereas the transition into the elongating state requires phosphorylation of the Serine 2 (Ser2P) by cyclin-dependent kinase 9 (CDK9) [8, 9, 10]. As shown previously, the Pol II Ser5P and Pol II Ser2P marks can be used to reliably visualize *in situ* the levels and location of recruited Pol II and elongating Pol II via immunofluorescence, respectively [27, 30]. We started our investigation using mouse embryonic stem cells (mESCs), a system in which the loss of Pol II clusters has been previously documented [23]. The mESCs were induced to exit pluripotency by growing them in RHB-A medium [37]. This protocol robustly commits mESCs to a rapid pluripotency exit towards neural fate [37, 50]. Images acquired by instant super resolution microscopy (iSIM) [51] revealed prominent clusters of recruited Pol II (Fig. 1A), which were previously seen in mESCs also by not phospho-specific visualization of Pol II [23]. In line with previous work, these clusters were lost at later differentiation timepoints [23], confirming their expected disappearance with the transition of mESCs from a pluripotent to a more differentiated state (Fig. 1B). Additionally, the signal strength for recruited Pol II (Pol II Ser5P) increased to a maximum at 12 h of differentiation and dropped again after 24 h (Fig. 1B). Signal strength for elongating Pol II (Pol II Ser2P) also reached a maximum at 12 h, but only fully decayed after 48 h (Fig. 1B).

**Fig. 1:**
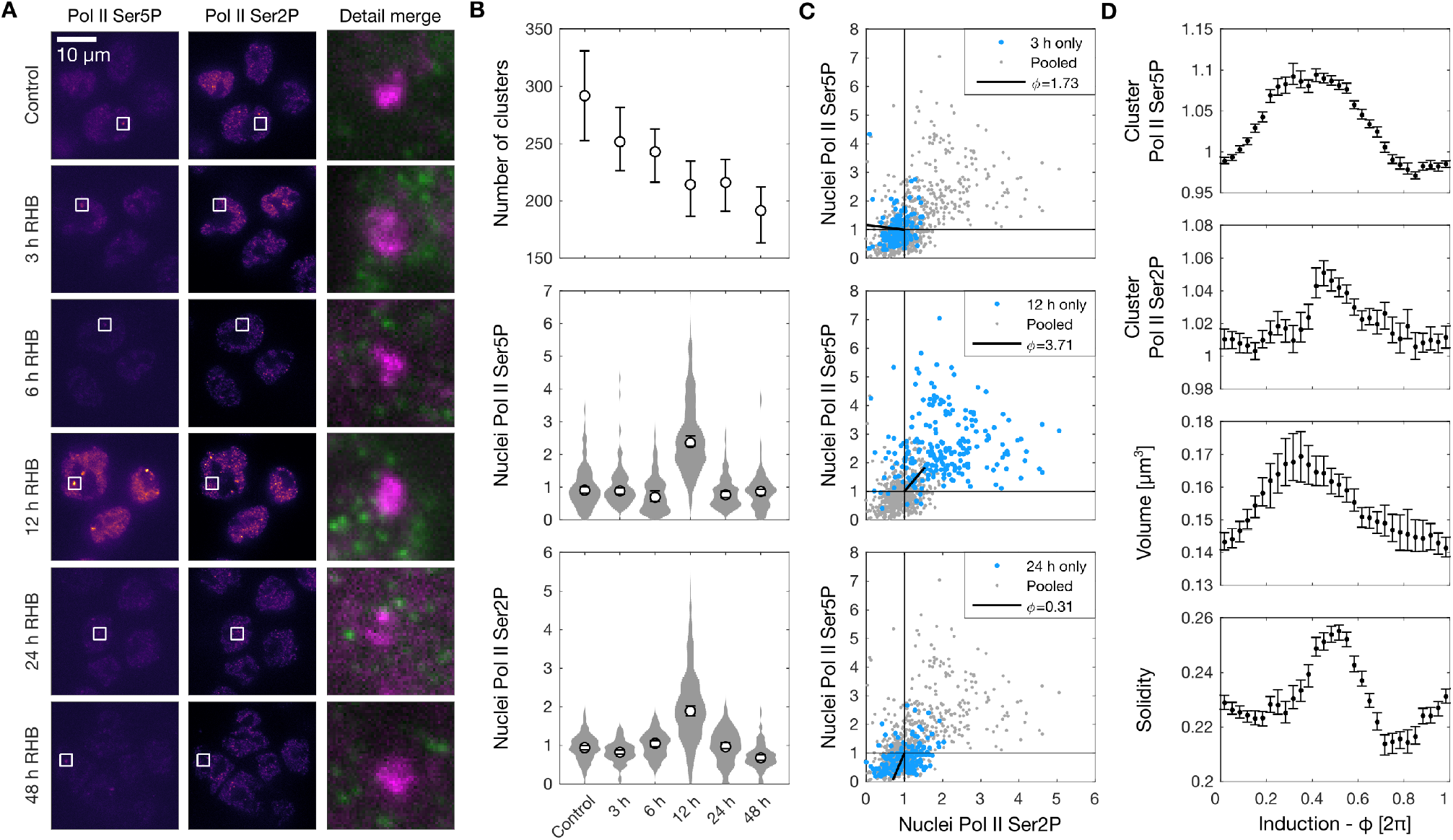
Formation and subsequent dispersal of RNA polymerase II (Pol II) clusters in mouse embryonic stem cells (mESCs) upon induced differentiation (using RHB-A medium, RHB). **A)** Example micrographs of mESC nuclei labeled against recruited Pol II (Ser5P) and elongating Pol II (Ser2P) by immunofluorescence. Single optical sections obtained by instant-SIM (iSIM) confocal microscopy. **B)** Quantification of whole-nucleus properties at different time points (0-48 h) during mESC differentiation. Number of Pol II Ser5P clusters per nucleus, minimal volume 0.03 *µ*m^3^ used as cut-off during analysis (mean with 95% bootstrap confidence intervals). Whole-nucleus immunofluorescence intensities normalized against the control condition (median with 95% confidence interval). Overall *n* = 119, 134, 111, 216, 162, 122 nuclei from two independent experimental repeats for every time point. **C)** Whole-nucleus Pol II Ser5P and Pol II Ser2P levels of all time points (including control treatment) are included into a joint scatter plot to define a phenomenological angular coordinate (*ϕ*) that represents progress in transcriptional induction. Whole-nucleus intensities from 3 h, 12 h, and 24 h time points are shown separately to illustrate differences in localization in the scatter plot, and how the angular coordinate *ϕ* can be used similar to a clock handle to represent differentiation progress. **D)** Nuclei are binned based on a sliding window for *ϕ*, and all Pol II Ser5P clusters from all nuclei in a given window are used to calculate the Pol II Ser5P and Pol II Ser2P intensity at these clusters, cluster volume, and solidity of the cluster (mean with 95% bootstrap confidence interval). A total of 17,219 clusters was included in this analysis.

Our microscopy data resolved Pol II Ser5P and Ser2P levels in single cells, so that the nucleus-wide levels of Pol II recruitment and elongation could be quantified (Fig. 1B). These levels exhibit a broad distribution, likely representing the fact that the exit from pluripotency in mESCs is not a highly synchronized process. Accordingly, we developed an approach to computationally sort cells according to levels of Pol II recruitment and elongation. Specifically, we introduced an angular coordinate (*ϕ*), which can be calculated using Pol II Ser5P and Ser2P intensity levels as coordinates in a scatter plot (Fig. 1C). The particular mathematical formulation we used to calculate *ϕ* ensures that Pol II recruitment was followed by the transition into elongation. Plotting of the mean *ϕ* for different points in the differentiation time course confirms that *ϕ* can capture the differentiation progress as intended (Fig. 1C).

Based on this angular coordinate, we advanced towards the analysis of Pol II state and shape of individual transcriptional clusters. Representative views of the Pol II foci are suggestive of systematic changes that occur with advancing differentiation (Fig. 1A). A comprehensive analysis supports this impression, showing that, indeed, levels of recruited Pol II transiently increase with progressing differentiation, followed by a transient increase in elongating Pol II (Fig. 1D). Our imaging analysis shows that Pol II clusters transiently increase in volume (Fig. 1D). We also analyzed the Pol II cluster solidity – a measure of how “rounded up” cluster shapes are – and we noticed that solidity first increases and then transiently drops, indicating a sequence where clusters first round up and then unfold (Fig. 1D). In other words, the clusters grow, round up, shrink in size again and finally disperse. To further consolidate the Pol II dynamics obtained for mESC differentiation, we repeated the analysis using cells in which exit from pluripotency was induced by withdrawal of leukaemia inhibitory factor (LIF) (SI Fig. 8). The results obtained for the two independent mESC differentiation protocols show that Pol II transcriptional clusters go through a reproducible sequence of changes in state and changes in shapes with progressing differentiation.

### A comparable formation-dispersal sequence is seen for prominent Pol II clusters during sperm cell precursor formation in a developed organ

To assess progressing differentiation of stem cells also in a developed organ instead of a laboratory cell culture system, we investigated the formation of fruit fly (*Drosophila melanogaster*) sperm precursor cells. These cells are formed by the differentiation of germ line stem cells, which are positioned in a hub at the apical tip of a testis (Fig. 2A). An increasing distance from the apical tip can be used as an indicator of the degree of differentiation, again providing an effective coordinate to sort cells by their degree of differentiation (Fig. 2B). In confocal microscopy images, prominent clusters of recruited Pol II can again be found, with different levels of recruited and elongating Pol II as well as different cluster shapes depending on the differentiation coordinate (Fig. 2B). A quantitative analysis of cells positioned along the differentiation coordinate, again, reveals a transient increase in levels of recruited Pol II, and a slower decrease in levels of elongating Pol II (Figure 2C). The area and the solidity of transcriptional clusters also undergo a transient increase (Fig. 2C). Taken together, we find that key features of the change in Pol II state and shape of transcriptional clusters match those observed during the differentiation of mESCs, indicating a conserved formation-dispersal sequence of Pol II clusters during stem cell differentiation also in the context of a developed organ.

**Fig. 2:**
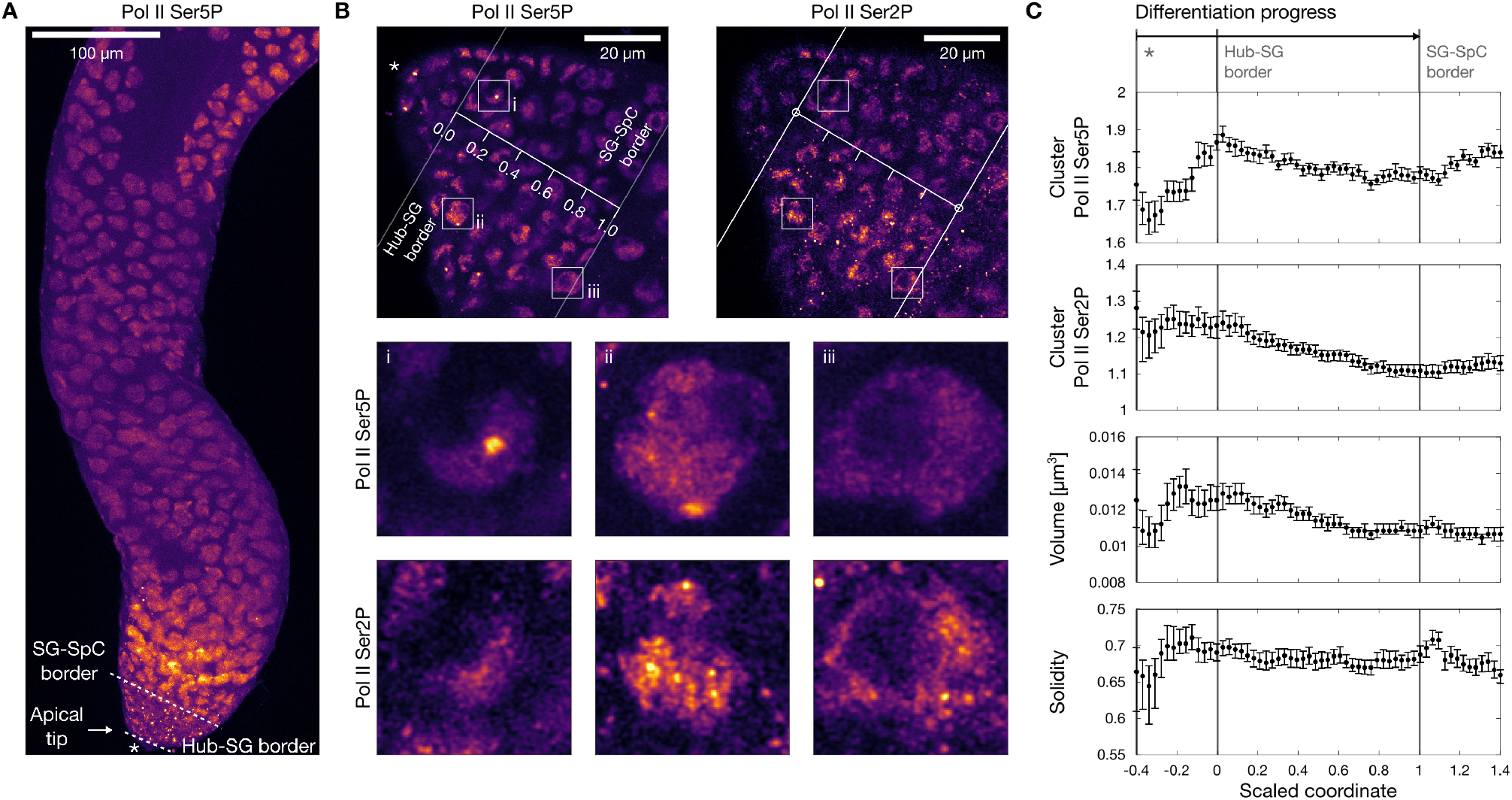
Formation and subsequent dispersal of RNA polymerase II (Pol II) clusters during sperm precursor formation in fruit fly testes. **A)** Representative confocal micrograph of a *Drosphila melanogaster* testis stained by indirect immunofluorescence against Pol II Ser5P and Ser2P. Apical tip, hub-spermatogonia (Hub-SG) border, and spermatogonia-spermatocyte (SG-SpC) border are indicated. **B)** Apical tip, the coordinate value 0.0 corresponds to the Hub-SG border and the value 1.0 corresponds to the SG-SpC border, providing a scaled differentiation coordinate (placed by visual assessment). **C)** Analysis of Pol II Ser5P cluster properties along the scaled coordinate. Median with 95% bootstrap confidence interval. For the solidity, a volume range selection of 0.01 to 0.035 *µ*m^3^ was applied to prevent biases from cluster volume differences. Overall, 60,114 clusters were analyzed from a total of 380 nuclei obtained from 10 samples from 3 independent experiments.

### Germ layer induction in a vertebrate embryo exhibits comparable formation and dispersal of transcriptional clusters in conjunction with reduction of active enhancer marks

As a third and final model of differentiation, we assessed the induction of stem cells towards the first germ layers during the natural development of zebrafish embryos. Not only does this transition occur as part of the natural development of a vertebrate organism, but also provides the ability of accurate collection of consecutive developmental stages. In particular, we collected early zebrafish embryos during the developmental time window ranging approximately from 3.7 to 5.3 hours post fertilization (hpf), thereby covering transcriptional activation of the zygotic genome (oblong and sphere stage) as well as the transition from pluripotency to persistent commitment to embryonic germ layers (dome, 30% epiboly, and 50% epiboly stage; Figure 3A) [52, 53, 54, 55, 56]. In all stages, prominent transcriptional clusters could be found, while also changes in the overall levels of recruited and elongating Pol II were visually apparent (Figure 3B). Immunofluorescence signal for recruited Pol II (Ser5P) was highest in the sphere stage, whereas the signal for elongating Pol II peaked later, in the 30% epiboly stage (Figure 3C). This sequence of a transient increase in recruited Pol II followed by a transient increase in elongating Pol II corresponds to the changes in Pol II state observed also for mESCs and in Drosophila testes. The number of prominent transcriptional clusters decreases with progressing development, mirroring the loss of clusters with progressing differentiation in mESCs (Figure 3C). Individual transcriptional clusters also exhibited a decrease in recruited Pol II, followed by a transient increase in elongating Pol II, as seen in the other experimental model systems (Figure 3C). Lastly, a transient increase in cluster volume as well as solidity could be seen, which is in line with the results from mESCs as well as Drosophila testes (Figure 3C). Based on this quantification, the formation and dispersal of prominent transcriptional clusters is characterized by similar changes in Pol II state and cluster shape in three experimental models of stem cell differentation: induction of mESCs towards a neural fate, formation of sperm precursor cells of Drosophila testes, and lineage induction in early zebrafish embryos.

**Fig. 3:**
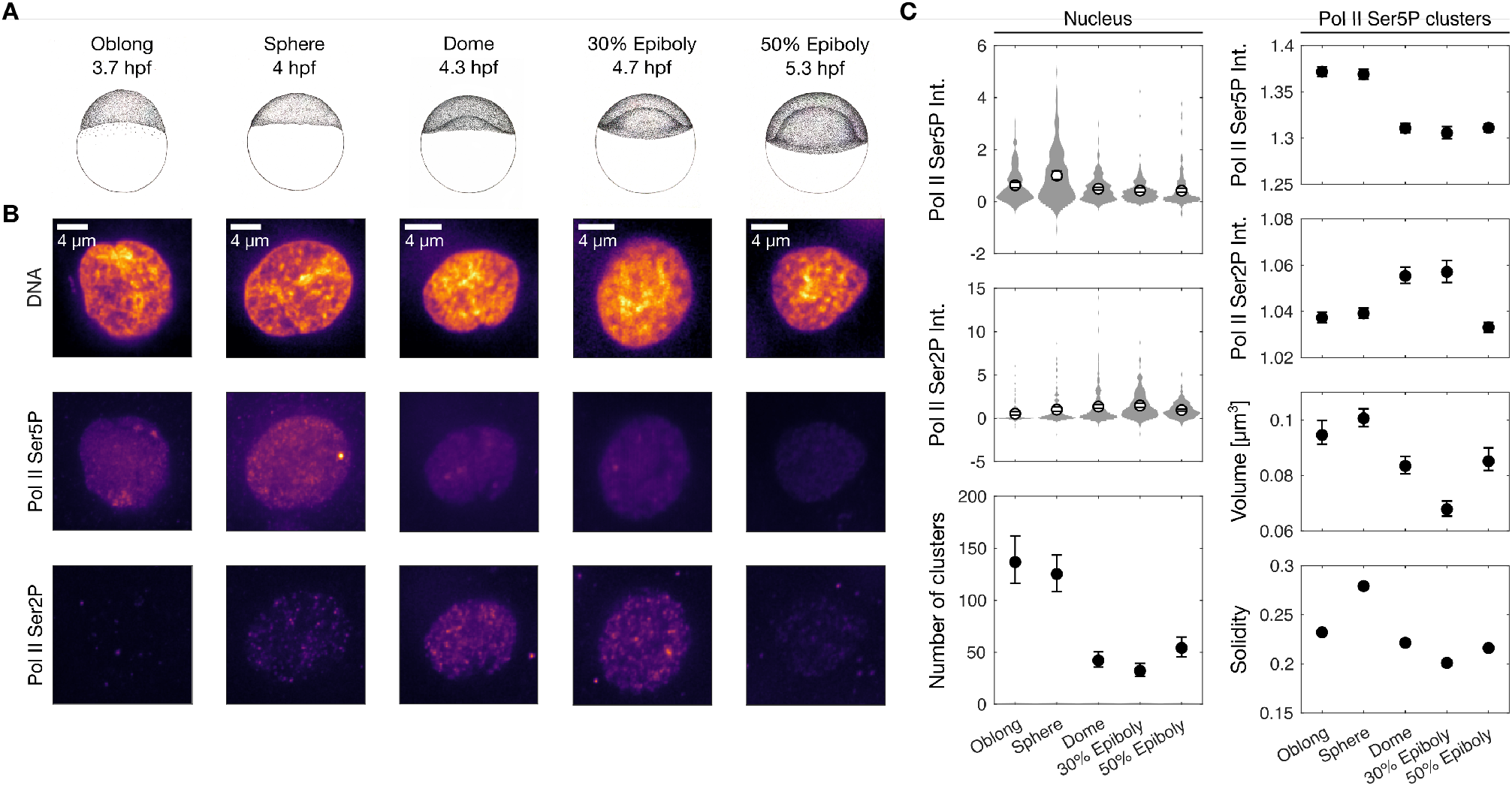
Formation and dispersal of RNA polymerase II (Pol II) clusters during germ layer induction in zebrafish embryos. **A)** Illustration of a zebrafish embryo approaching full transcriptional activation (oblong stage), reaching pluripotency (sphere stage), and undergoing progressive gastrulation (dome, 30% epiboly, and 50% epiboly stages). The embryo is seen laterally, with the animal cap containing the blastula cell mass on top. Hours of development post fertilization (hpf) are indicated for each stage. **B)** Representative micrographs of nuclei of zebrafish embryos in the developmental stages displayed in panel A. DNA and Pol II were fluorescently labeled and nuclear mid-section images were acquired by instant-SIM confocal microscopy. DNA was labeled with Hoechst 33342; immunofluorescence was used to detect recruited RNA polymerase II (Pol II Ser5P) and elongating RNA polymerase II (Pol II Ser2P). **C)** Quantification of fluorescence intensity in whole nuclei (*n* =173, 211, 291, 138, 162, normalized against sphere stage) and at Pol II clusters (*n* =5,568, 11,427, 12,285, 18,876, 20,298, intensities normalized against the median of each analyzed nucleus), data obtained from *N* = 6, 4, 6, 4, 4 embryos from three independent experimental repeats. For the analysis of clusters, a minimal volume of 0.03 *µ*m^3^ was required for clusters to be included in the analysis. Data are shown as mean with 95% bootstrap confidence intervals.

Epigenetic states, including prominently the H3K27ac mark, were found to change rapidly during development, affecting chromatin organization and transcriptional activity of genes [57, 58, 59, 60]. High levels of the H3K27ac mark are commonly associated with active enhancers and Pol II recruitment in stem cells [61, 62]. Additionally, a partial reduction of superenhancer-associated H3K27ac levels has been previously documented for progressing differentiation in mammalian embryonic stem cells (mESCs) [63, 64]. In early zebrafish embryos, the H3K27ac mark has been shown to precede and support the formation of prominent transcriptional clusters [65, 28, 66, 30, 67] and to be markedly reduced by the completion of gastrulation [68]. Considering the accuracy of developmental staging of zebrafish embryos, we applied immunofluorescence to assess changes in the H3K27ac mark during the progressing differentiation process (Fig. 4A). We observe that the H3K27ac level remains unchanged throughout the first three stages (oblong to dome, Fig. 4B), followed by a drop in the last two stages (30% epiboly and 50% epiboly). Considering that the epiboly stages of development are associated with the onset of morphogenesis and locking-in of cell lineages [52, 53], our observations imply that the exit from embryonic pluripotency is indeed associated with a reduction of H3K27ac-marked regulatory regions that can support cluster formation. Overall, our experiments indicate a conserved transition during the differentiation of stem cells. In all three experimental model systems, a transient increase in recruited Pol II is followed by a later transient increase in elongating Pol II, along with growth and rounding up of Pol II clusters, followed by their dispersal with further differentiation. In the developing zebrafish embryo, we could thus confirm the reduction of active enhancer marks with progressing differentiation, as previously observed for differentiation in mESCs.

**Fig. 4:**
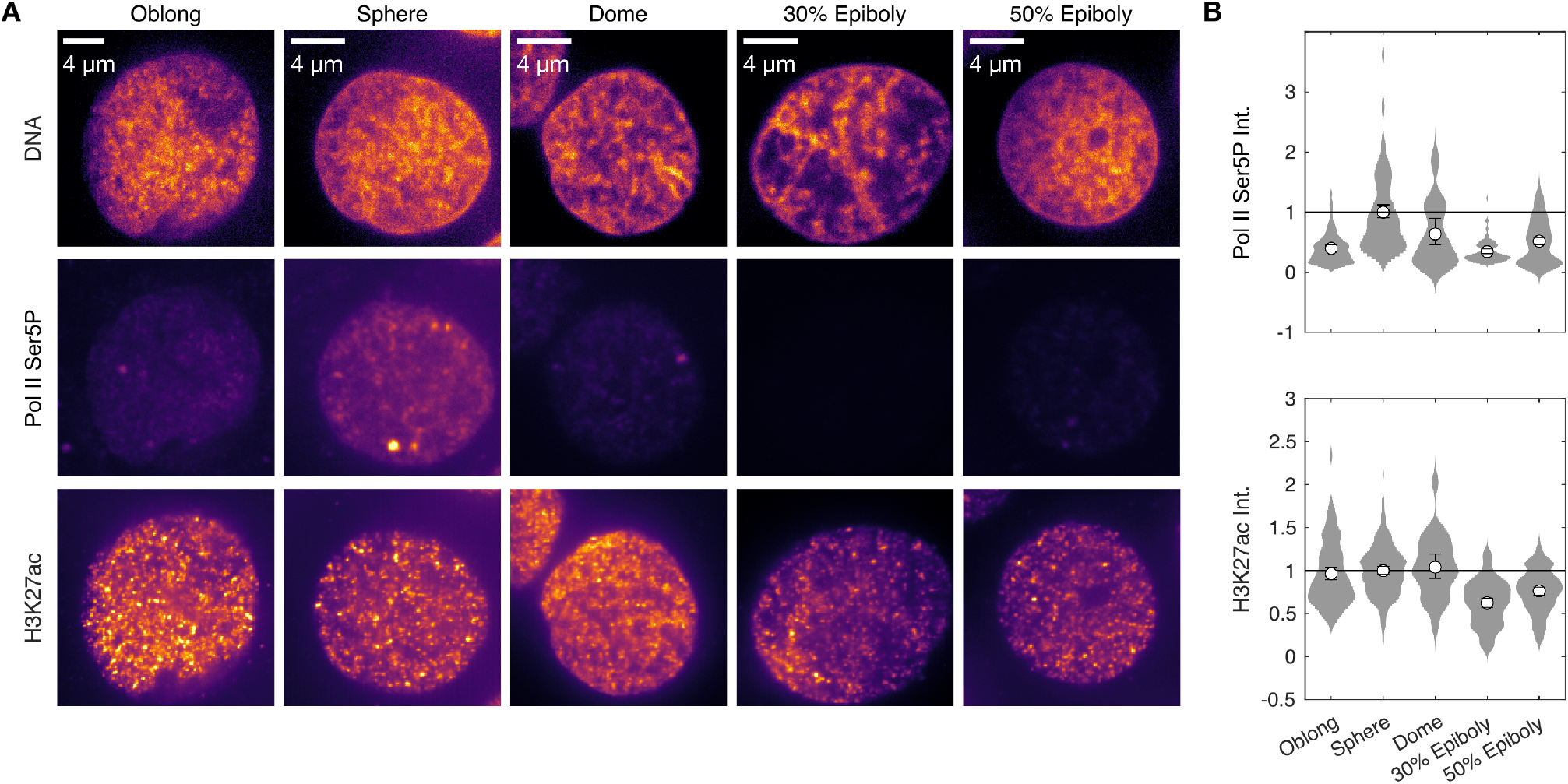
Histone acetylation marks of active enhancers are reduced in late stages of germ layer induction. **A)** Representative nuclear mid-sections of nuclei in fixed sphere-stage zebrafish embryos, recruited RNA polymerase II (Pol II Ser5P) and lysine-23-acetylated histone 3 (H3K27ac) were stained by indirect immunofluorescence, DNA with Hoechst 33342. **B)** Quantification of Pol II Ser5P and H3K27ac intensities. Overall *n* = 81, 117, 27, 99, 165 nuclei were recorded from 4, 4, 2, 4, 3 embryos from a single immunofluorescence repeat to obtain more comparable fluorescence intensities. Mean with 95% bootstrap confidence interval, normalized against values obtained at the sphere stage.

### A surface condensation model reproduces the experimentally observed formation and dispersal of transcriptional clusters

Considering the similarity of the observations in all three experimental models of differentiation, we theorized that cluster formation and dispersal proceed on the basis of a general underlying biophysical mechanism. Accordingly, we aimed to provide a generalized theoretical model for the formation of transcriptional clusters. Our requirement of such a model was that the experimentally determined, nucleus-wide levels of available condensation surface (H3K27ac), recruited Pol II (Ser5P), and transcriptional activity (Pol II Ser2P) are sufficient to serve as inputs to the model, so that simulations would produce cluster morphologies (Figure 5A). Previous work proposed a surface condensation model of transcriptional cluster formation, which can accept H3K27ac, Pol II Ser5P, and Pol II Ser2P levels as input parameters [30]. In particular, H3K27ac-marked super-enhancers facilitate the condensation of a liquid phase that contains recruited Pol II, whereas increased levels of elongating Pol II induce the unfolding or even dispersal of transcriptional condensates formed in this fashion (Figure 5B). These aspects of the model imply that the reduction in super-enhancers that are available as condensation surfaces as well as increased transcription levels seen in our experiments could reduce the overall number of prominent transcriptional clusters.

**Fig. 5:**
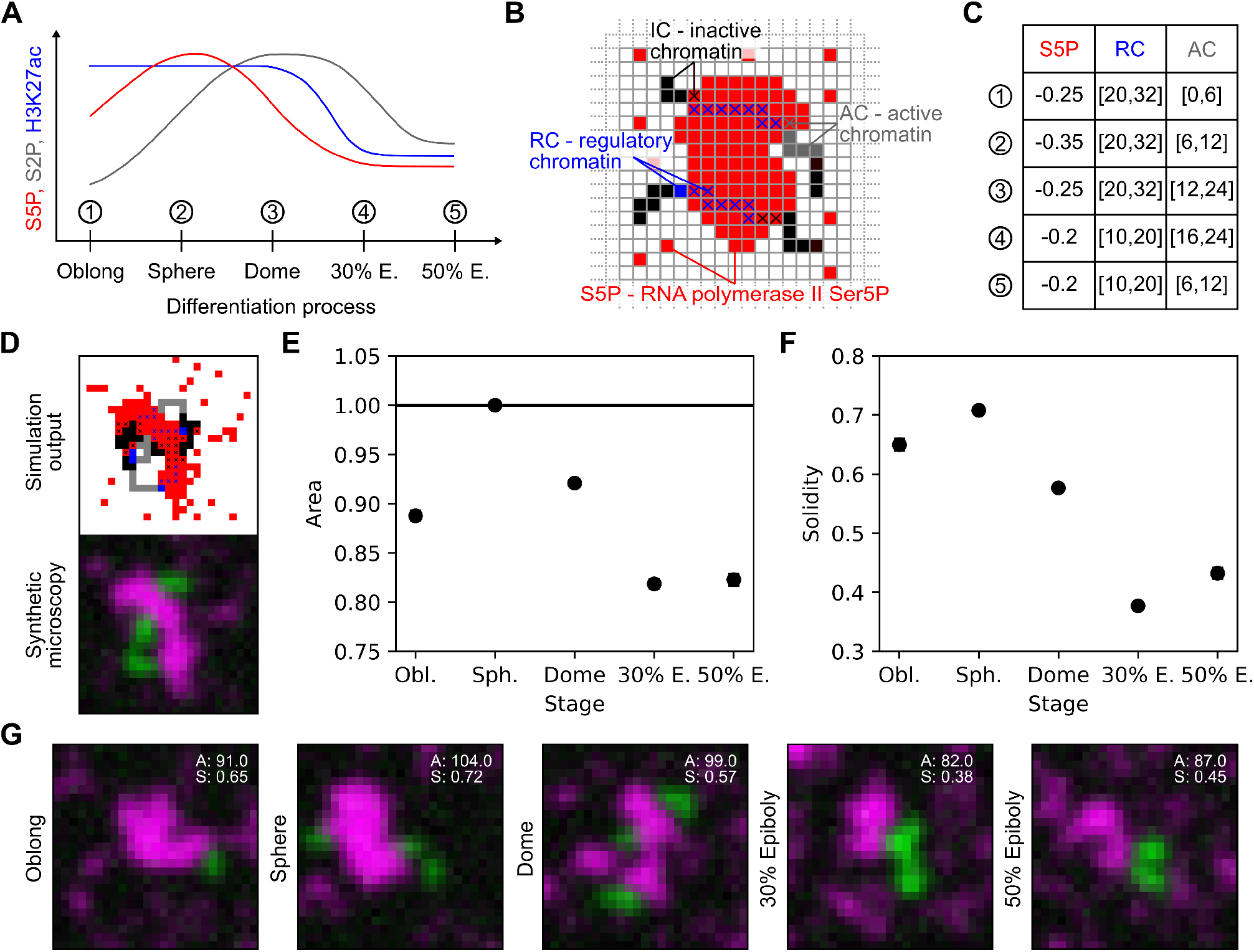
Simulations of surface condensation reproduce changes in the cluster area and solidity observed throughout the differentiation process. **A)** Sketch of experimental results (zebrafish embryos) that summarizes the changes in levels of recruited Pol II (S5P), transcription activity (S2P) and available surface (H3K27ac). **B)** Lattice representation of surface condensation model, consisting of polymers with three subregions (IC - inactive chromatin, RC - regulatory chromatin, AC - active chromatin) and liquid material that can form clusters and contains recruited RNA Pol II (S5P). **C)** Selection windows of simulation parameters representing the different stages. **D)** Simulation output and conversion to synthetic microscopy images. **E)** Resulting cluster area as mean with 95% bootstrap confidence interval. Normalized by mean value of sphere stage. **F)** Resulting cluster solidity as mean with 95% confidence bootstrap confidence interval. **G)** Synthetic microscopy example images for each stage, area (*A*) and solidity (*S*) as indicated.

For numerical simulations of the theoretical model, we adapted the coarse-grained lattice kinetic Monte-Carlo (LKMC) approach [69] from previous work on surface condensation (Figure 5B) [30]. With a proper choice of parameters, this model can describe the formation of surface-supported condensates in the sub-saturated regime, where bulk phase separation does not occur. The model relies on generic interaction laws, and thus delivers a first-principle theoretical prediction of the overall system dynamics. The lattice simulations contain single particles that represent the material forming the clusters enriched in recruited Pol II, which exhibit self-affinity (S5P, red, interaction energy *w_S_*_5_*_P−S_*_5_*_P_ <* 0). Chromatin is represented by block copolymer chains with different subregions: regions of “inactive” chromatin (IC, black, self affinity *w_IC−IC_ <* 0), followed by regions of regulatory chromatin (RC, blue) that have an affinity for the red species (*w_RC−S_*_5_*_P_ <* 0) and thereby can serve as a condensation surface, and directly adjacent regions of active chromatin (AC, gray) that is repelled by red particles (*w_AC−S_*_5_*_P_ >* 0). We performed simulations containing polymer chains with randomized lengths of blue and gray block regions for different affinities between red particles, which all fall within the sub-saturated regime with respect to the red particles [30], thereby obtaining a total of 300 different lattice simulation runs (for more details, see Material and Methods). To select those simulations that represent the five developmental stages of zebrafish embryos, we defined corresponding selection windows that can be applied to the randomly assigned polymer assemblies (see Fig. 5C). Lastly, to compare simulation outputs to the microscopy data, we produced synthetic microscopy images by adding a Gaussian blur filter approximating limited microscope resolution and detector noise (Poisson noise) to the lattice model output (see Fig. 5D). We then extracted cluster area and solidity with an image analysis pipeline that corresponds to the analysis of our actual microscopy images (Fig. 5E, F). As was the case in our experiments, the area as well as the solidity of clusters transiently increased at the sphere stage, and dropped to the lowest values at the 30% epiboly and 50% epiboly stages (Fig. 5E, F). To supplement our quantitative analysis, we also provide representative synthetic microscopy images for each stage, which compare favorably with clusters seen in actual microscopy data (Fig. 5G). Taken together, simulations based on condensation facilitated by super-enhancers as a condensation surfaces reproduced key features of the formation and dispersal of transcriptional clusters as observed in our experiments.

### Experimental perturbations of super-enhancer status and transcription elongation induce theoretically explicable changes in cluster morphology

Our findings suggest that a reduction in the levels of H3K27ac-marked super-enhancers and an increase in transcriptional activity can explain the loss of prominent transcriptional clusters during differentiation. To test both parameters as causative, we proceeded to chemically perturb super-enhancer efficacy with the BET domain inhibitor JQ-1, which affects the status of H3K27ac-marked enhancers [70, 65] (30-min treatment), and with the CDK9 inhibitor flavopiridol, which hinders the transition of Pol II into elongation (30-min treatment), in zebrafish embryos collected at the sphere stage [71] (Figure, 6A, for full validation of inhibitor effects see SI Fig. 9). JQ-1 treatment led to a reduction in cluster volume, and a small but statistically significant reduction in cluster solidity (Figure 6B). In contrast, flavopiridol treatment led to an increase in cluster solidity (Figure 6B). To approximate the inhibitor treatments in our simulations, we reduced the affinity to the condensation surface (JQ-1) or removed transcriptionally active chromatin (flavopiridol) (Figure 6C). The resulting synthetic microscopy images reproduce the experimentally detected changes in cluster area and solidity for JQ-1 as well as flavopiridol treatment (Fig. 6D). In conclusion, super-enhancer status as well as transcriptional activity causally contribute to the morphology of transcriptional clusters, in line with the detailed predictions from the surface condensation model of cluster formation.

**Fig. 6:**
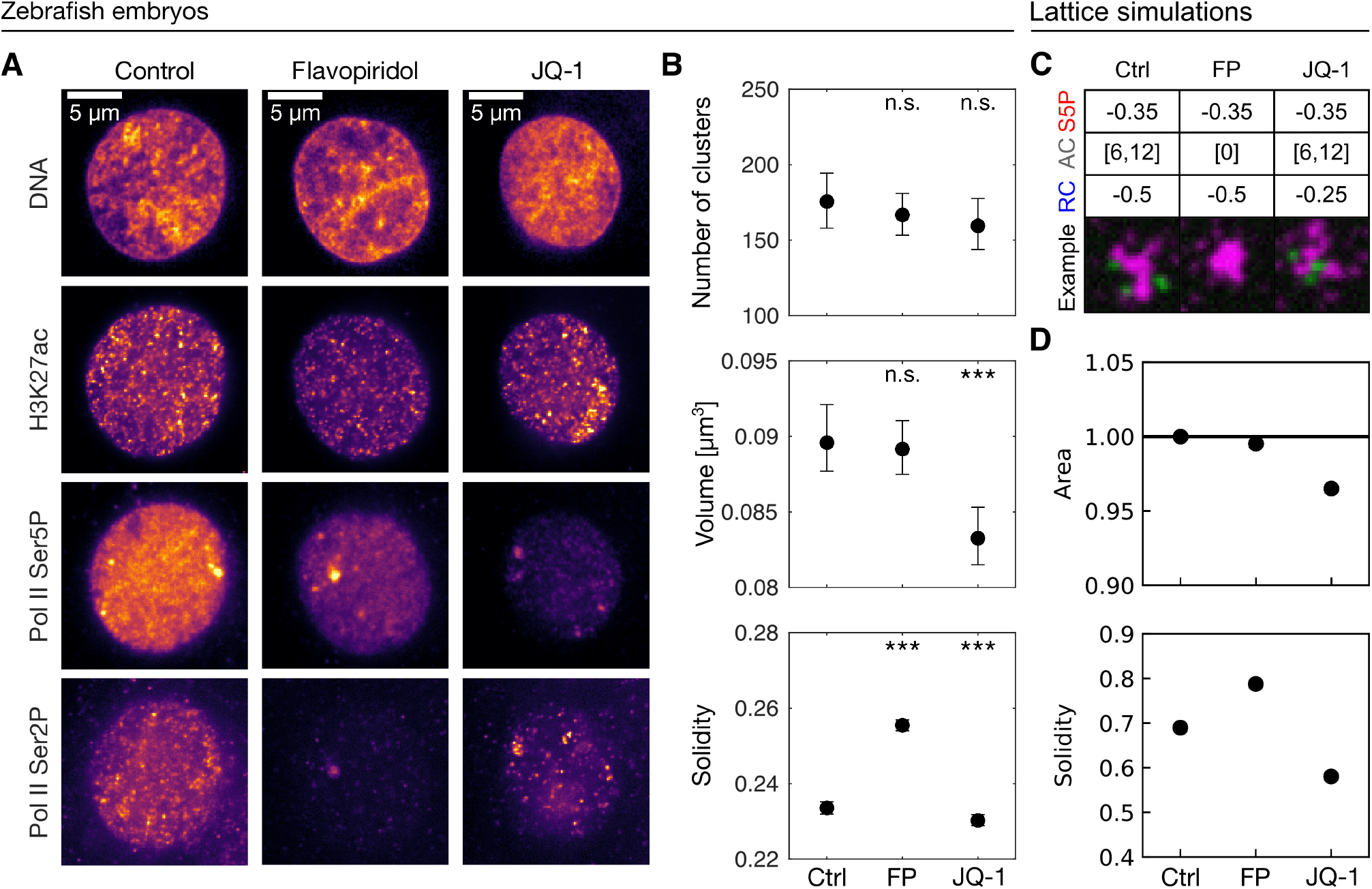
Effects on transcriptional clusters caused by chemical inhibitor treatment of pluripotent zebrafish embryos. **A)** Example micrographs of nuclear mid-sections; lysine-23-acetylated histone 3 (H3K27ac), recruited RNA polymerase II (Pol II Ser5P), and elongating RNA polymerase II (Pol II Ser2P) were stained by indirect immunofluorescence, DNA with Hoechst 33342. **B)** Quantification of number of clusters, cluster area and cluster solidity for control (Ctrl, sphere stage), flavopiridol (FP) and JQ-1 treatment. P-values of two-tailed permutation test of differences of the mean relative to the control treatment: 0.9033, 0.4102; 1.5260, 0.0002; 0.0002, 0.0030; Bonferronicorrected for multiple comparison. n=193, 292, 251 nuclei for comparison of number of clusters; n=35,292, 52,017, 39,864 Pol II Ser5P clusters for comparison of volume and solidity (minimum volume for solidity calculation 0.03 *µ*m^3^). Mean with 95% bootstrap confidence interval, *** indicates statistical significance with *P <* 0.001, n.s. indicates no statistical significance with *P ≥* 0.05. **C)** Parameter changes and example images for simulated treatments. **D)** Quantification of cluster area and cluster solidity from simulations of the three experimental conditions, Mean with 95% confidence interval (n=50,50,50 simulations). Statistics of simulated control and treatments for cluster area and solidity.

## Discussion

In this study, we found that the dispersal of prominent transcriptional clusters during differentiation occurs via a sequence of changes in the regulatory state and the shape of clusters that is conserved across different experimental models of differentiation. Based on our simulations of cluster formation via surface condensation, this conserved sequence can be seen as a journey through a cluster shape space defined along three dimensions: (i) level of recruited Pol II, (ii) available condensation surface provided by super-enhancers, and (iii) transcriptional activity (Fig. 7). For any “position” in this shape space, a particular size and shape of transcriptional clusters can be expected on the basis of our simulations of surface condensation. During the process of differentiation, a typical trajectory through this shape space is traversed, where an initial increase in Pol II recruitment is followed by a transiently increased level of transcriptional activity and finally a partial loss of enhancers that serve as condensation surfaces. The growth and subsequent dispersal of transcriptional clusters implied by this shape space trajectory directly reflect our experimental observations for progressing differentiation. In summary, our findings mark a surface condensation model as a coherent explanation for the highly conserved changes in the spatial organization of transcription with ongoing differentiation of stem cells.

**Fig. 7:**
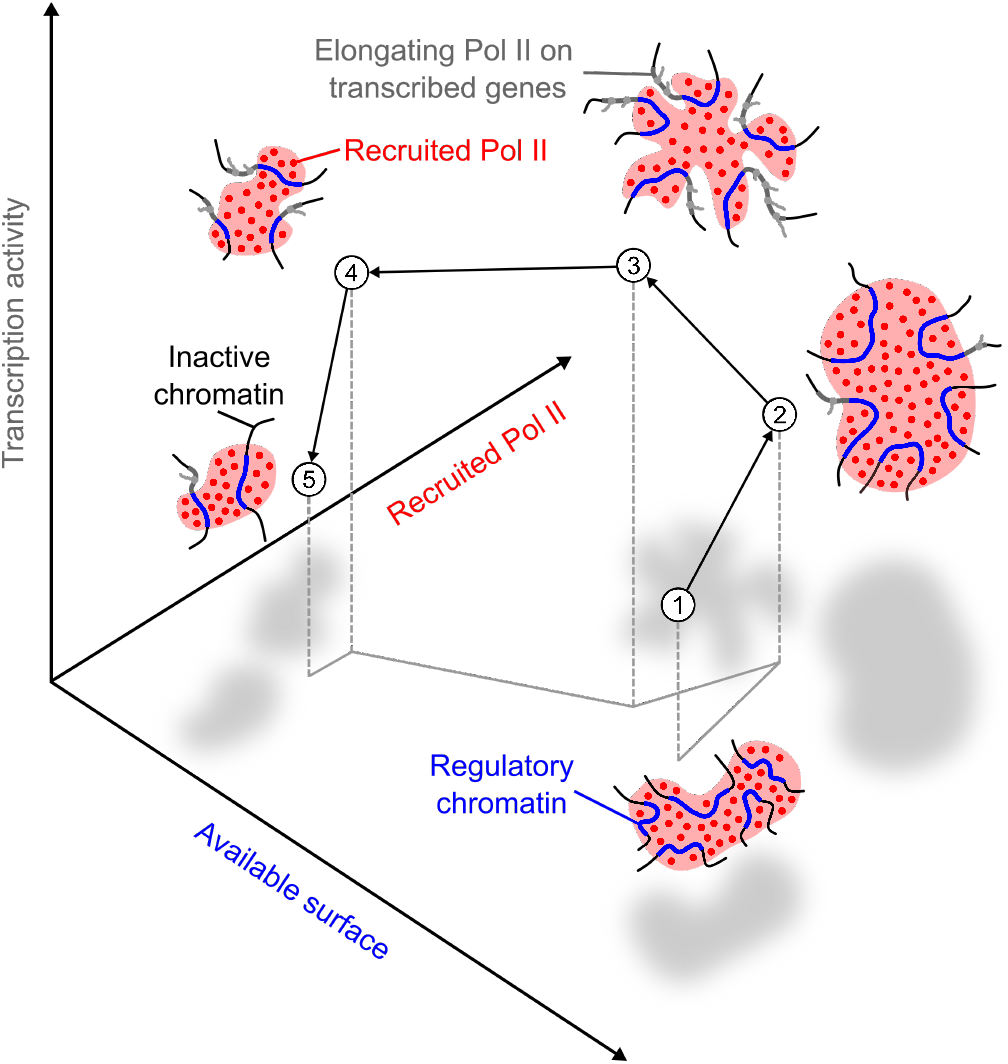
Diagram explaining the concept of a cluster shape space. A position within the shape space is defined by the level of recruited Pol II, available condensation surface, and transcription activity. Over the course of differentiation, a stereotyped trajectory is traversed in this space, and clusters appear in different sizes and shapes depending on their position in this space.

Originally described as “transcription factories” [12, 14], transcriptional clusters have been investigated intensely over the past years [2, 3]. Transcriptional clusters are widely observed, including in human cells [12, 13, 18, 16, 72, 6, 19, 20, 21], mouse erythroblasts and mESCs [22, 23], Drosophila embryos [24, 25, 26], and zebrafish embryos [30, 31, 32]. Accordingly, the process by which these clusters form has been actively discussed [73, 4, 1, 15]. Several studies have suggested a role for liquid-liquid phase separation in the formation of Pol II clusters, but also revealed that *in vivo* concentrations typically fall below the saturation concentration required for the formation of phase-separated droplets [38, 23, 39, 8, 40, 41, 42, 43]. One mechanism that allows the formation of condensates at sub-saturated concentrations is provided by surface condensation that is mediated by DNA strands with transcription factor binding motifs [44, 45]. Here, the size and location of condensates is controlled by the presence of relevant binding sites that act as a binding surface. A similar contribution to condensate formation by super-enhancer regions has been previously observed [38, 40]. Further, the formation of Pol II clusters in zebrafish embryos could be reproduced by simulations of surface condensation, where regulatory chromatin acts as a condensation surface [30]. Previous work found that, once formed, transcriptional clusters can also exhibit an internal spatial organization, including compartments at the scale of *∼*100 nm that correspond to consecutive steps of transcription regulation [9, 10, 21, 74]. Separation of Pol II recruitment and transcript elongation, for example, could be explained in terms of the exclusion of transcribed genes and their RNA transcripts from a liquid phase enriched in recruited Pol II [11, 75]. Here, transcribed genes take on the role of an effective amphiphile, which can unfold or even split apart transcriptional clusters [27, 30, 31]. A comparable dispersal of DNA-nanomotif condensates by synthetic amphiphile particles has been demonstrated in a fully artificial, cell-free model system [76]. In our study, a simulation based on surface condensation and the dispersing effect of transcription reproduced the step-wise changes in transcriptional clusters through the differentiation process. These results support a surface condensation model of *in vivo* transcription cluster formation, where further reshaping of clusters occurs in connection to increased transcriptional activity.

While our work provides an explanation of how the conserved formation and dispersal of transcriptional clusters can be achieved, the question remains how such clusters might contribute to the particular transcriptional programs associated with stem cells and their differentiation [77, 78]. Work over the recent years has contributed to a model of gene-regulatory hubs, where multiple enhancers come into contact and provide a localized context that can be visited by genes, which thereby undergo transcriptional regulation [79, 80, 81, 82, 83]. In embryonic development, persistent enhancer hubs were found to be visited by different genes depending on differentiation and transcriptional activation [84, 85]. According transient interactions of genes with enhancers that control their transcription has been seen in live imaging [86, 87] as well as pseudo-time reconstruction from fixed cells [31]. Hubs involving enhancers and genes also form prior to transcriptional activation, with an involvement of recruited Pol II, seemingly providing a configuration that is poised for subsequent transcriptional activation during differentiation processes ([88, 89, 37]). In line with this idea of preparation prior to transcriptional activation, an increase in Pol II recruitment [90, 91] as well as a preemptive formation of gene-gene and gene-enhancer loops [92, 88] was found. Additionally, super-enhancers underlying long-range contacts as well as gene expression changes are partially deactivated upon exit from pluripotency and activated upon induction of new cell types, supporting a model where transcriptional hubs would be formed and subsequently dispersed via the activation and deactivation of super-enhancers [79, 93, 63, 64]. While the overall structure of transcriptional hubs is not changing with changes in transcription, more nuanced changes can be seen in a number of example systems. In mice, specific enhancer-promoter interactions increase with differentiation into different tissue types [94]. Also in mice, neural differentiation leads to changes specifically in promoter-enhancer contacts *in vitro* [93] and the organization of long genes *in vivo* [95]. In zebrafish embryos, super-enhancers become increasingly compacted [96] and contribute to long-range contacts [97] with developmental progress into and beyond embryonic pluripotency. In Drosophila, long-range contacts are equally established prior to and independently of transcriptional activation, via clustering of the pluripotency-associated transcription factor Zelda [98, 99, 100]. Similar clustering of pluripotency-associated transcription factors and chromatin modifiers into “transcription bodies” was seen also in zebrafish [29, 32] as well as in mESCs [23]. Transcription factor binding sites are clustered at regions where cohesin accumulates, indicating that similar transcription factor clustering might occur in human cells [101]. In conclusion, the spatial clustering of enhancers and transcription-associated factors emerges as a mechanism by which stem cells might prepare for subsequent differentiation into specific cell types. The condensation and subsequent dispersal of transcriptional clusters seen in our work appears as one aspect of this preparation for and subsequent execution of stem cell differentiation. Considering that this transition is conserved across several experimental models of differentiation and can be traced back to generic biophyiscal mechanisms, a similar condensation-dispersal sequence of transcriptional clusters would also be expected in cases of stem cell differentiation beyond the systems assessed in this study.

## Materials and Methods

### Mouse embryonic stem cell culture

#### Cell culture and differentiation

Mouse cells were grown at 37°C in a 5% (v/v) CO_2_ incubator as described in [37]. In brief, mESCs (ESC-46C; [102]) were grown in GMEM medium (Invitrogen), supplemented with 10% (v/v) Fetal Calf Serum (FCS; BioScience LifeSciences), 2,000 U/ml LIF (Millipore), 0.1 mM *β*-mercaptoethanol (Invitrogen), 2 mM L-glutamine (Invitrogen), 1 mM sodium pyruvate (Invitrogen), 1% penicillin-streptomycin (Invitrogen), 1% MEM Non-Essential Amino Acids (Invitrogen) on gelatin-coated (0.1% v/v) Nunc T25 flasks. The medium was changed every day, and cells were split every other day. Before sample collection (control time point), mESCs were plated on gelatin-coated (0.1% v/v) Nunc 10-cm dishes in serum-free ESGRO Complete Clonal Grade Medium (Millipore) to which was added 1,000 U/ml LIF. Medium was changed to the cells every 24 h and cells were collected at 48 h. mESC batches were tested for mycoplasma infection. Early neural differentiation was carried out as described in [37]. mESCs were plated with high density (1.5 × 10^5^ cells/cm^2^) in serum-free ESGRO Complete Clonal Grade Medium (Millipore) to which 1,000 U/ml LIF was added. After 24 h, mESCs were washed 3 times with PBS (without magnesium and calcium), incubated in PBS for 3 min at room temperature, and then dissociated by incubating in 0.05% (v/v) Trypsin (Gibco) for 2 min at 37*^◦^*C. mESCs were plated onto 0.1% (v/v) gelatin-coated 10-cm dishes (Nunc) at 1.6*×*10^6^ cells/dish in RHB-A (Takara-Clontech), changing media every 24 h. These differentiating cells were collected after 3 h, 6 h, 12 h, 24 h, and 48 h in RHB-A. Alternatively, for the LIF withdrawal experiment, mESCs were plated onto 0.1% (v/v) gelatin-coated 10-cm dishes (Nunc) at 1.6*×*10^6^ cells/dish in GMEM medium (see above) without LIF. Differentiating cells were collected after 3 h, 6 h, 12 h, 24 h, and 48 h of LIF withdrawal. Upon harvesting, cell suspension was fixed with 2% PFA for 30 min at room temperature and then centrifuged (5 min, 320 g). To increase the mechanical stability of cells, the supernatant was removed and a secondary fixation step was carried out by applying 8% formaldehyde in PBS for 30 min at room temperature, followed by centrifugation (5 min, 320 g) and removal of the supernatant.

#### Immunofluorescence

For all steps that include replacement of liquids, sample tubes were centrifuged (1 min, 800g) before liquid removal. To begin immunostaining, cells were fixed again with a higher concentration of formaldehyde (8% FA in PBST, 20 min, room temperature), then washed three times with PBST. Cells were permeabilized with 0.5% Triton X-100 in PBS (10 min, room temperature), then washed three times with PBST. Cells were blocked with 4% BSA in PBST (30 min, room temperature). Cells were incubated in primary antibody mix (1:1,000 rabbit anti-Pol II Ser2P; 1:1,000 rat anti-Pol II Ser5P in 4% BSA in PBST, Tab. 1) overnight at 4*^◦^*C, then washed three times with PBST. Cells were incubated in secondary antibody mix (4% BSA in PBST with 1:1,000 anti-rat IgG Alexa 594 and 1:1,000 anti-rabbit IgG Alexa 488, Tab. 1) overnight at 4*^◦^*C, then washed three times with PBST. Samples were post-fixed with 8% formaldehyde in PBST for 15 min at room temperature, then washed three times with PBST. For sample mounting, liquid was removed as far as possible without removing cells, and replaced by VectaShield H-1000 with 2 *µ*M Hoechst 33342 (1:10,000 dilution from 20 mM stock). Cells were resuspended and transferred to 8-chamber microscopy slides (ibidi *µ*-slide, glass bottom #1.5 selected cover glass) using a P200 micropipette (100 *µ*l per chamber).

**Tab. 1:** List of primary and secondary antibodies used for imunofluorescence. All primary antibodies were validated in a previous publication [30].

| Target | Type | Clone/Conjugate | Supplier | Cat. No. |
| --- | --- | --- | --- | --- |
| **Primary antibodies** |  |  |  |  |
| Pol II S5P | Mouse IgG | 4H8 | Abcam | ab5408 |
| Pol II S5P | Rat IgG | 3E8 | Active motif | 61986 |
| Pol II S2P | Mouse IgM | H5 | Biolegend | 920204 |
| Pol II S2P | Rabbit IgG | EPR18855 | Abcam | ab193468 |
| H3K27ac | Rabbit IgG | EP16602 | Abcam | ab177178 |
| **Secondary antibodies** |  |  |  |  |
| Anti-mouse IgM | Goat | Alexa 594 | Invitrogen | A21044 |
| Anti-rabbit | Donkey | Alexa 488 | Invitrogen | A21206 |
| Anti-rabbit | Goat | Alexa 647 | Thermofisher | A21244 |
| Anti-rabbit | Goat | Alexa 488 | Thermofisher | A11006 |
| Anti-rat | Goat | Alexa 594 | Invitrogen | A11007 |
| Anti-rat | Goat | Alexa 488 | Thermofisher | A11066 |
| Anti-rat | Goat | Alexa 647 | Thermofisher | A21247 |

#### Microscopy

The prepared samples were recorded using a VisiTech iSIM high-speed superresolution confocal microscope based on the instant-SIM principle [51], built on a Nikon Ti2-E stand. A Nikon 100x Oil Immersion objective (NA 1.49, SR HO Apo TIRF 100xAC Oil) and excitation lasers at 405, 488, 561 and 640 nm were used. Two Hamamatsu ORCA Flash4.0 V3 cameras were used for dual color acquisition on the basis of a 561-nm long-pass beam splitter. In a given experimental repeat, the illumination and acquisition setting were kept constant.

#### Image analysis

Nuclei were segmented by the Pol II Ser5P intensity applying a Gaussian blur (kernel width *σ* = 1.0 *µ*m) for the foreground and for background subtraction (*σ* = 10 *µ*m). To exclude edge effects, nuclei masks were eroded with a disk of diameter 0.5 *µ*m, followed by a 3D hole-filling topological operation. Only nuclei of minimal volume of 10 *µ*m^3^ and minimal solidity 0.8 were retained for further analysis. A ring-shaped cytoplasm mask was formed by extending from the nuclear mask, with a ring beginning at a distance of 1.0 *µ*m from the nuclear mask, reaching to an outer distance of 1.5 *µ*m. For cluster detection, Gaussian blur was applied for segmentation of the Pol II Ser5P channel (*σ* = 0.02 *µ*m), with background removal (Gaussian blur with *σ* = 0.1 *µ*m). Segmentation of clusters was done with the robust background threshold method (1.5 standard deviations above mean of all pixel intensities). Individual objects were joined into clusters using DBSCAN with *ϵ* = 0.5 *µ*m. Clusters below a minimal volume of 0.03 *µ*m^3^ were removed from further analysis. Fluorescence intensities inside clusters were mean intensities averaged over all pixels in the cluster segmentation mask and normalized against the median intensities over the entire nucleus that a given cluster resides in. Nuclei were sorted according to an angular coordinate, which was calculated as

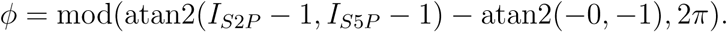

By this formulation, *ϕ* = 0 corresponds to a point where the Pol II Ser5P signal is minimal, which can be seen as prior to the increased formation of Pol II Ser5P clusters.

Microscopy data and image analysis scripts are available via Zenodo repository: https://doi.org/10.5281/zenodo.8013242

### Drosophila testes

#### Drosophila testes sample preparation

Prior to sample preparation, a 96-well plate with conical wells was prepared as followed: 4 wells were filled for each of the following solutions: PBS, 4% formaldehyde in PBS, 0.1% Triton X-100 in PBS, PBS with 0.1% Tween-20 (PBST) and 1% BSA in PBST. Flies were moved from a chosen vial into a new one. The old vial, containing pupae, but not adult flies, was incubated for 1 h at room temperature to obtain freshly hatched flies. Subsequently, the flies were anaesthetized using CO_2_ and transferred to a fly pad. The male flies were collected and transferred into a 500 *µ*l drop of PBS on a dissection dish (black background). The flies were dissected to obtain testes, and 2-3 pairs of testes were gently placed into the first well filled with PBS. They were moved through four wells with PBS. Next, they were moved through all four wells with 4% formaldehyde to assure equilibration of the formaldehyde concentration, finally allowing for 20 min of fixation in the last formaldehyde well. Further, the testes were permeabilized by moving the samples through all four wells filled with 0.1% Triton X-100 and an incubation step of 30 min in the last well with Triton X-100. Lastly the samples were moved through the four wells filled with PBST and washed by being left in the last one for 15 min.

#### Immunofluorescence

Immunostaining was performed directly after permeabilization. To block the samples for non-specific antibody binding, the samples were moved through four wells of a 96-well culture plate filled with 1% BSA in PBST and left to incubate for 45 min at room temperature in the last well. During this blocking procedure, a primary antibody solution (1:300 rabbit anti-Pol II Ser2P and 1:300 rat anti-Pol II Ser5P in 1% BSA in PBST, Tab. 1) was prepared and two remaining wells of the 96-well plate were filled. The testes were transferred through the two wells of primary antibody solution and incubated in the second well in a moist dark chamber overnight at 4*^◦^*C. The samples were moved through four wells with PBST and incubated in the last one for 15 min. The secondary antibodies solution (1:300 goat anti-rabbit conjugated to Alexa 647 and 1:300 goat anti-rat conjugated to Alexa 488 in 1% BSA in PBST, Tab. 1) was prepared and two wells were filled. The testes were moved through both wells with the secondary antibodies solution and incubated in the second well in a moist dark chamber over night at 4*^◦^*C. The samples were then moved through four wells with PBST for washing and incubated in the last one for 15 min. Finally, the testes were mounted on a poly-L-lysine slide in 30 *µ*l Vectashield with 1:10,000 Hoechst 33342, covered with a #1.5 cover slip on top and sealed with nail polish.

#### Microscopy

Microscopy data from *Drosophila melanogaster* testes were recorded using an LSM 900 confocal fluorescence microscope with Airyscan 2 with a Plan-Apochromat 63x/1.40 Oil DIC M27 objective. In a given experimental repeat, the illumination and acquisition setting were kept constant.

#### Image analysis

Nuclei and Pol II clusters were segmented based on the Pol II Ser5P channel by image processing steps that are identical to those used for mESCs. Parameter values were adjusted: nuclei segmentation foreground Gaussian kernel *σ* = 0.6 *µ*m, nuclei background subtraction Gaussian kernel *σ* = 8 *µ*m, nuclear segmentation mask erosion 0.1 *µ*m, topological hole-filling in 2D, nuclei minimal volume 10 *µ*m^3^ and minimal solidity 0.4; cytplasmic mask reaching 0.5 *−* 1.5 *µ*m, cluster segmentation foreground Gaussian blur *σ* = 0.001 *µ*m, cluster segmentation background subtraction Gaussian blur *σ* = 0.1 *µ*m, robust background removal threshold 2 standard deviations above mean. DBSCAN *ϵ* = 0.3 *µ*m, minimal cluster volume 0.005 *µ*m^3^.

To assign the segmented nuclei a coordinate that represents differentiation progress, the Hub-SG border and the SG-SpC border were marked manually with a point coordinate in each recorded image stack. Based on these two points, an axis could be digitally drawn into the image, allowing the projection of all the centroid *xy*-positions of each nucleus to be projected to this axis. A projected value of 0 corresponds to the beginning of the differentiation process, a projected value of 1 to the completion of the differentiation process.

Microscopy data and image analysis scripts are available via Zenodo repository: https://doi.org/10.5281/zenodo.8014328

### Zebrafish embryos

#### Zebrafish husbandry

All zebrafish husbandry was performed in accordance with the EU directive 2010/63/EU and German animal protection standards (Tierschutzgesetz §11, Abs. 1, No. 1) and is under supervision of the government of Baden-Württemberg, Regierungspräsidium Karlsruhe, Germany (Aktenzeichen35-9185.64/BH KIT). Embryos used for the different experiments were obtained through spontaneous mating of adult zebrafish. Collected embryos were dechorionated with Pronase, washed three times with E3 embryo medium, once with 0.3x Danieau’s solution, and subsequently kept in agarose-coated Petri dishes or 6-well plates in 0.3x Danieau’s solution at 28.5*^o^*C.

#### Sample preparation

When zebrafish embryos reached chosen stages (oblong, sphere, dome, 30% epiboly and 50% epiboly), five embryos each were transferred into 2 ml Eppendorf tubes filled with 1 ml embryo fixative (0.3x Danieu’s, 2% formaldehyde (FA), 0.2% Tween-20). For fixation the samples were left at 4*^◦^*C overnight. On the next day, the samples were washed three times with PBST for 5 min. Afterwards the embryos were transferred into a Petri dish filled with PBST and the animal caps of the embryos were carefully separated from the yolk by using two fine forceps. The animal caps were transferred back into a tube and either stored at 4*^◦^*C or were directly used. The animal caps were permeabilized with 1 ml permeabilization mix (PBS with 0.5% Triton X-100) for precisely 15 min at room temperature. The samples were washed three times with PBST for 5 min. Finally the animal caps were directly used for immunostaining.

#### Whole Zebrafish embryo inhibitor treatment

Zebrafish embryos were treated with the chemical inhibitors Flavopiridol (stock concentration 12.5 mM) or JQ-1 (stock concentration 50 mM). In both cases, treatment proceeded for 30 min, starting such that 30 min of treatment finished when reaching sphere stage. Control embryos developed in parallel and were equally collected in sphere stage.

#### Immunofluorescence

Directly after permeabilisation the immunostaining was carried out. The samples were blocked with 4% BSA in PBST for 30 min at room temperature. Depending on the experimental setup, primary antibodies (anti-H3K27ac, anti-Pol II Ser2P and anti-Pol II Ser5P in 4% BSA in PBST) were applied overnight at 4*^◦^*C. The samples were washed three times with PBST for 5 min. The samples were washed once with 4% BSA in PBST. Depending on the experiment, different secondary antibodies were used (for details on antibodies, see Tab. 1). The secondary antibodies were applied in 4% BSA in PBST overnight at 4*^◦^*C. The samples were washed three times with PBST for 5 min. For the two-color immunofluorescence of Pol II Ser5P and Pol II Ser2P, a previously established and validated antibody combination was used (primary antobodies: 1:300 rabbit anti-Pol II Ser2P, ab193468; 1:300 rat anti-Pol II Ser5P in 4% BSA in PBST; secondary antibodies: 1:300 anti-rat IgG Alexa 594 and 1:300 anti-rabbit IgG Alexa 488 in 4% BSA in PBST) [30]. PBST was carefully removed from the animal caps of the embryos under the microscope to prevent destroying and loosing of the animal caps. For mounting 30 *µ*l of Vectashield H-1000 with 5 *µ*M Hoechst 33342 (1:2,500 dilution from a 20 mM stock concentration) was added to every sample. The animal caps were transferred with a P20 micropipette on a microscope slide. The slide was covered with a cover slip (#1.5, selected) and the slides were sealed with nail polish.

For the three-color immunofluorescence of Pol II Ser5P, Pol II Ser2P, and H3K27ac, a mouse IgM and a rat IgG primary antibody had to be used, introducing a small risk of crosstalk in the secondary antibody staining step. To reduce this risk of crosstalk, two rounds of indirect immunofluorescence were carried out. First, a mouse IgM primary antibody (1:300 anti-Pol II Ser2P in 4% BSA-PBST, see Tab. 1)) was incubated overnight at 4*^◦^*, then an anti-mouse IgM secondary antibody (1:300, anti-mouse IgM, conjugated to Alexa 594). Subsequently, a rat IgG and a rabbit IgG primary antibody (1:300 rabbit anti-H3K27ac and 1:300 rat anti-Pol II Ser5P) were incubated overnight, followed by overnight incubation of according secondary antibodies (1:300 anti-rabbit conjugated to Alexa 488 and 1:300 anti-rat conjugated to Alexa 647). The potential crosstalk from the mouse IgM-based detection into the rat IgG-based detection was not apparent (Fig. 10). Raw data and example images for crosstalk assessment are available via the following Zenodo repository: https://doi.org/10.5281/zenodo.8031698

#### Microscopy

Microscopy data from Zebrafish embryos were recorded using a VisiTech iSIM high-speed super-resolution confocal microscope as described for the recording from mESCs.

#### Image analysis

The segmentation of nuclei and Pol II clusters was based on the DNA and Pol II Ser5P channels, respectively, and was implemented in a pipeline with steps identical to the image processing for mESCs and sperm precursor cells. For the assessment of Pol II clusters in different stages (Fig. 3), parameter values were adjusted: nuclei segmentation foreground Gaussian kernel *σ* = 3 *µ*m, nuclei background subtraction Gaussian kernel *σ* = 10 *µ*m, nuclear segmentation mask erosion 1.5 *µ*m, topological hole-filling in 3D, nuclei minimal volume 10 *µ*m^3^ and minimal solidity 0.8; cytoplasmic mask reaching 1.0 *−* 1.5 *µ*m, cluster segmentation foreground Gaussian blur *σ* = 0.001 *µ*m, cluster segmentation background subtraction Gaussian blur *σ* = 0.1 *µ*m, robust background removal threshold 1.5 standard deviations above mean. DBSCAN *ϵ* = 0.65 *µ*m, minimal cluster volume 0.03 *µ*m^3^. Additional sorting steps were not necessary because embryos were sorted by developmental stage. The raw image data and analysis scripts are provided via the following Zenodo repository: https://doi.org/10.5281/zenodo.8013305

For the assessment of H3K27ac levels at Pol II clusters in different stages (Fig. 4), Pol II Ser5P signal was used for segmentation of nuclei as well as Pol II clusters. Parameter values were adjusted: nuclei segmentation foreground Gaussian kernel *σ* = 3.0 *µ*m, nuclei background subtraction Gaussian kernel *σ* = 10 *µ*m, nuclear segmentation mask erosion 1.5 *µ*m, topological hole-filling in 3D, nuclei minimal volume 10 *µ*m^3^ and minimal solidity 0.8; cytoplasmic mask reaching 1.0 *−* 1.5 *µ*m, cluster segmentation foreground Gaussian blur *σ* = 0.001 *µ*m, cluster segmentation background subtraction Gaussian blur *σ* = 0.1 *µ*m, robust background removal threshold 1.5 standard deviations above mean. DBSCAN *ϵ* = 0.65 *µ*m, minimal cluster volume 0.005 *µ*m^3^. Additional sorting steps were not necessary because embryos were sorted by developmental stage. The raw image data and analysis scripts are provided via the following Zenodo repository: https://doi.org/10.5281/zenodo.8019735 For the assessment of Pol II clusters upon flavopiridol and JQ1 treatment (Fig. 6), nuclei were segmented based on DNA signal and Pol II clusters based on Pol II Ser5P signal. Parameter values were adjusted: nuclei segmentation foreground Gaussian kernel *σ* = 3 *µ*m, nuclei background subtraction Gaussian kernel *σ* = 10 *µ*m, nuclear segmentation mask erosion 1.5 *µ*m, topological hole-filling in 3D, nuclei minimal volume 10 *µ*m^3^ and minimal solidity 0.8; cytoplasmic mask reaching 1.0 *−* 1.5 *µ*m, cluster segmentation foreground Gaussian blur *σ* = 0.001 *µ*m, cluster segmentation background subtraction Gaussian blur *σ* = 0.1 *µ*m, robust background removal threshold 1.5 standard deviations above mean. DBSCAN *ϵ* = 0.65 *µ*m, minimal cluster volume 0.005 *µ*m^3^. The raw image data and analysis scripts are provided via the following Zenodo repository: https://doi.org/10.5281/zenodo.8019764

### Biophysical lattice model for cluster description

For the theoretical biophysical description of the experimentally observed phenomena, we use our existing lattice kinetic Monte-Carlo (LKMC) model of surface condensation [30]. While all parts that are essential to reproduce our results are described in here, the mentioned publication and publicly hosted simulation scripts can be used as reference for a more detailed description of the model.

#### Model components

The model consists of four different species (see SI Fig.11A). First, liquid material containing recruited RNA Pol II (S5P, red), the species that actually forms clusters, is represented by single lattice sites. In addition, a polymer that serves as surface for cluster formation is modeled as a chain of connected lattice sites. To represent different activities and functions of chromatin, the polymer consists of three different subregions, lined up in the following order: inactive chromatin (IC, black), regulatory chromatin (RC, blue) and active chromatin (AC, gray), and flanked again by inactive chromatin.

#### Inter- and intraspecies affinities

To enable formation of clusters, self affinity (*w_S_*_5_*_P−S_*_5_*_P_ <* 0) is assumed (see SI Fig.11B). The parameter is adjusted so that canonical phase separation does not happen. This is the case for all chosen values *w_S_*_5_*_P−S_*_5_*_P_ ∈* [*−*0.2*, −*0.25*, −*0.35]. The different polymer subregions have different affinities to S5P and with themselves. IC is subject to self affinity (*w_IC−IC_ <* 0), reflecting a tendency for compaction, and is neutral to other chromatin species and S5P. RC has no self affinity, but instead affinity to S5P (*w_RC−S_*_5_*_P_ <* 0). AC and S5P repel each other (*w_AC−S_*_5_*_P_ >* 0), as it was shown that elongating Pol II is excluded from transcriptional clusters.

#### Move set

S5P is implemented as single particles within the lattice and can move to all eight nearest neighbor lattice sites (see SI Fig.11C). Movements are only prohibited in the case an S5P particle is located at the lattice border, since movements outside the lattice boundaries are not permitted, or in the case the targeted neighboring site is already occupied by another S5P particle. For the polymer, the known Verdier-Stockmayer move set, consisting of kink-jump, end-bond flip and crankshaft move, is used [103, 104, 69]. This restricted move set allows simulation of self-avoiding polymers. S5P and polymers have no space exclusion interactions and evolve on different lattice layers.

#### Initial configuration

All simulations are performed within a lattice of same size (25 *×* 25 lattice sites) (see SI Fig.11D). The numbers of S5P (*N_S_*_5_*_P_* = 100) and polymers (*N_Polymer_* = 4) are kept constant for all simulations. The single S5P particles are randomly distributed within the lattice at the beginning of a given simulation. Polymers overall have the same length (*L_Polymer_* = 20), whereas the length of each subregion (*N_IC_*, *N_RC_* and *N_AC_*) is varied for different developmental stages or treatments. *N_RC_ ∈* [0, 2, 4, 6, 8] (per polymer), *N_AC_ ∈* [0, 3, 6] (per polymer), *N_IC_* as much as necessary to fill up to the total length *L_Polymer_*. At the beginning of the simulation, four polymer chains are stacked vertically in the center of the lattice and their orientation is alternated.

#### Simulation framework

The coarse-grained LKMC simulations [105, 69] act as a Gillespie-type algorithm [106] and run rejection-free (see SI Fig.11E). After setting the initial configuration at the beginning of a given simulation, all lattice sites are checked for possible system transitions. These transitions are then saved together with their rates *k* (determined by energy difference and move type) and the total system energy *k_total_* as sum of all transition rates. Out of this catalog, one transition is chosen by the tower sampling method, which takes into account a random number *r ∈* [0, 1] and the system energy. This transition is then performed, immediately followed by a local update of the affected positions and proceeding to choosing the next transition. The choosing and performing loop is repeated *N* times. Most simulations were performed with a total of 1 *×* 10^6^ LKMC iteration steps.

#### Simulation analysis

To be able to compare the simulation output to experimental results and use the same image analysis pipeline, synthetic microscopy images are produced (see SI Fig.11F). To this end, we blur the images produced by the lattice model with Gaussian kernel (*σ* = 1) and add Poisson-distributed random numbers (*λ* = 5, resulting ranomd numbers divided by 100) to simulate detector noise. For cluster detection, blurred distributions of S5P are used, to which a global segmentation threshold of 0.35 is applied, cluster labeling (connected-component labeling, CCL) is performed, and labeled clusters are retained if they exceed a minimal area of 10. We used ergodic sampling (every 1 *×* 10^4^ step) for simulation snapshots after 8 *×* 10^5^ initial time steps for equilibration. For each of these sampling time points, cluster area, solidity, and S5P and S2P intensities are measured. The numerical simulations as well as the image analysis are carried out using Python (Spyder).

Data sets and scripts for theoretical model are available via the following Zenodo repository: https://doi.org/10.5281/zenodo.7966557

## Acknowledgements

This work was funded via the Helmholtz program Natural, Articifical, and Cognitive Information Processing (NACIP). TK, IW, VZ, and LH received funding from the priority program 2191 of the Deutsche Forschungsgemeinschaft (SPP2191). CF was supported by the clinical research unit 5002 of the Deutsche Forschungsgemeinschaft (KFO5002). SE and LH received seed funding of the KIT center for Materials in Technical and Life Sciences (MaTeLiS). We thank Hiroshi Kimura for discussions and Xenia Tschurikow for illustrations of zebrafish embryo developmental stages as well as polymerase clusters.

## Supplementary Material

**Fig. 8:**
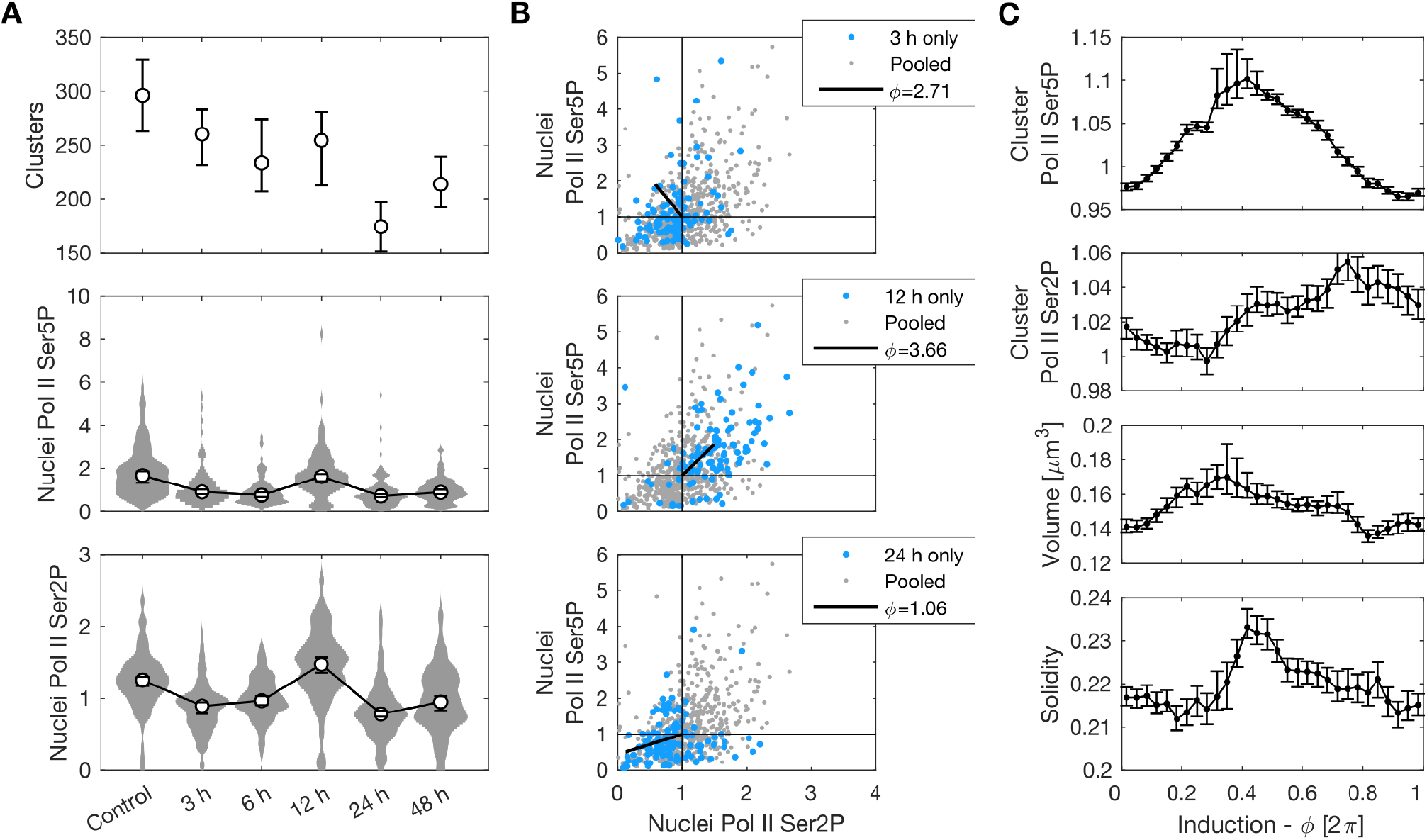
Formation and dispersal of RNA polymerase II (Pol II) clusters in mouse embryonic stem cells upon leukaemia inhibitory factor (LIF) withdrawal. **A)** Changes in whole-nucleus properties at different time points after LIF withdrawal. Number of clusters per nucleus considers Pol II Ser5P clusters with a minimal volume of 0.03 *µ*m^3^ (mean with 95% bootstrap confidence interval). Whole-nucleus intensities of recruited Pol II (Pol II Ser5P) and elongating Pol II (Pol II Ser2P) are normalized against control (median with 95% bootstrap confidence interval). *n* = 112, 108, 101, 110, 135, 135 nuclei from 2 independent experimental repeats. **B)** Whole-nucleus Pol II Ser5P and Pol II Ser2P levels of all time points (including control treatment) are included into a joint scatter plot to define a phenomenological angular coordinate (*ϕ*) that represents progress in transcriptional induction. Whole-nucleus intensities from 3 h, 12 h, and 24 h after LIF withdrawal are shown separately to illustrate differences in localization in the scatter plot, and how the angular coordinate *ϕ* can be used similar to a clock handle to represent differentiation progress. **C)** Quantification of properties of large Pol II Ser5P clusters in a sliding window analysis (mean with 95% bootstrap confidence interval). A minimal volume cut-off filter (0.08 *µ*m^3^) was applied. A total of 13,525 clusters was included in this analysis.

**Fig. 9:**
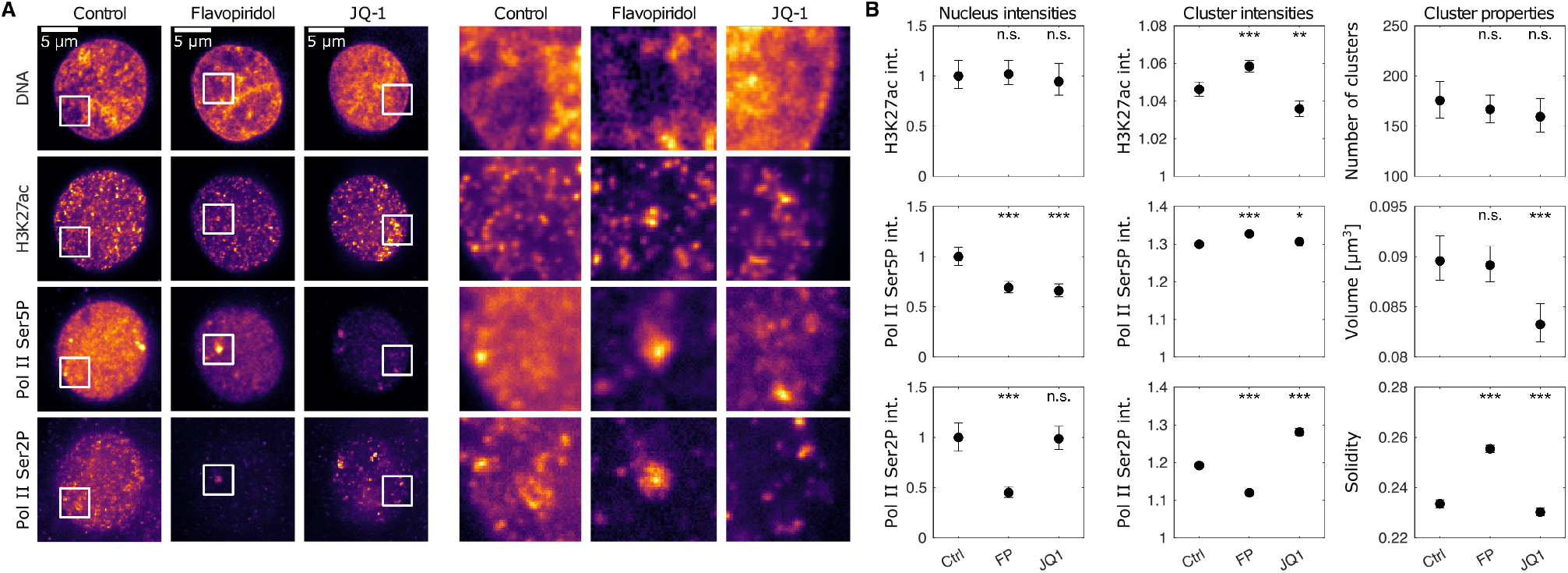
Full analysis – effects on transcriptional clusters caused by chemical inhibitor treatment of pluripotent zebrafish embryos. **A)** Example micrographs, nuclear mid-sections, look-up tables kept the same for all immunofluorescence channels. **B)** Quantifications for control (Ctrl, sphere stage), flavopiridol (FP) and JQ-1 treatment. P-values of two-tailed permutation test of differences of the mean relative to the control treatment: 1.6798, 1.2147; 0.0002, 0.0002; 0.0002, 1.7916; 0.0002, 0.0018; 0.0002, 0.0102; 0.0002, 0.0002; 0.9033, 0.4102; 1.5260, 0.0002; 0.0002, 0.0030; Bonferroni-corrected for multiple comparison. n=193, 292, 251 nuclei for comparison of nucleus intensities as well as number of clusters; n=35,292, 52,017, 39,864 Pol II Ser5P clusters for comparison of cluster intensities, volume, and solidity (minimum volume for solidity calculation 0.03 *µ*m^3^; mean with 95% bootstrap confidence interval).

**Fig. 10:**
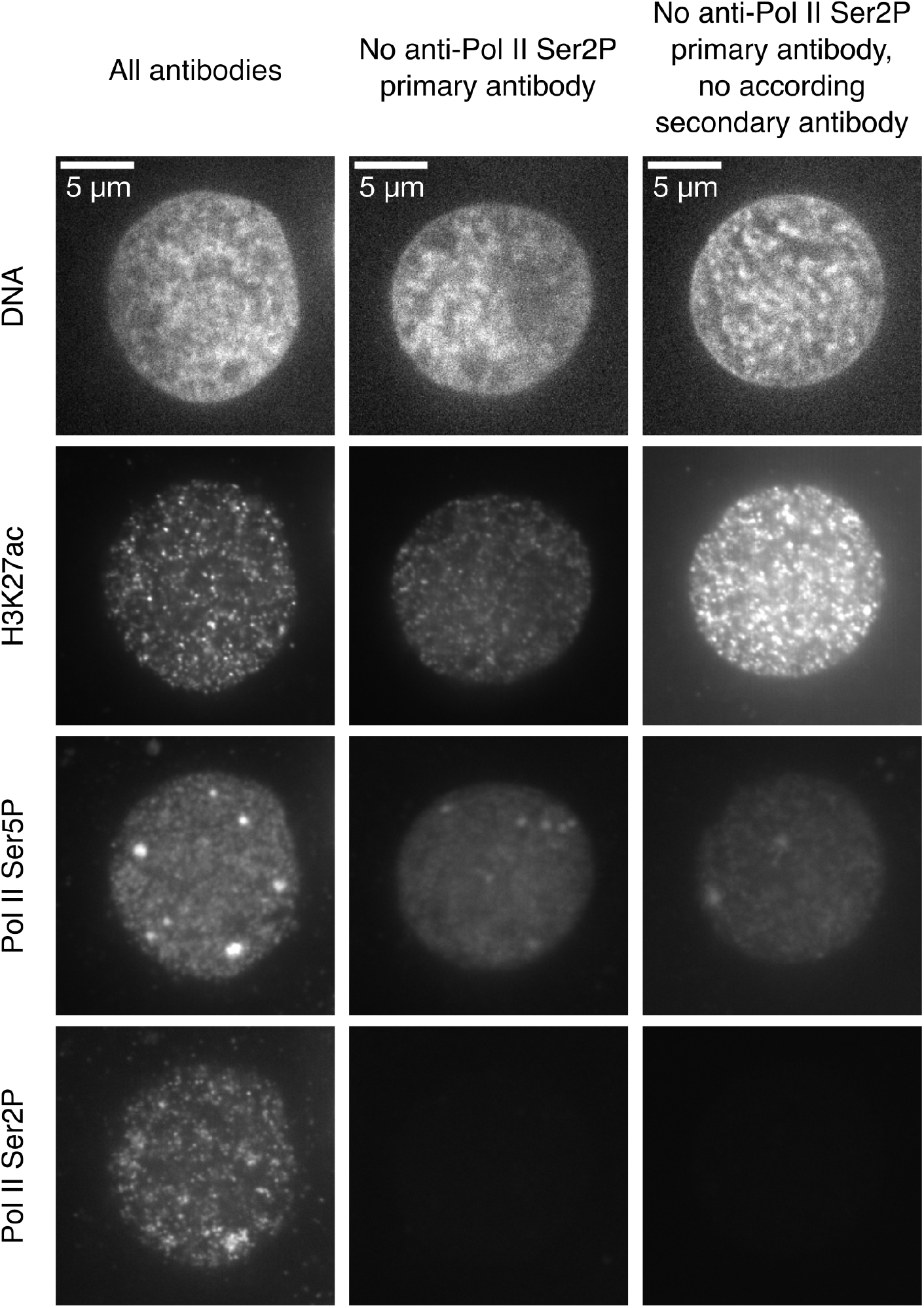
Detection of elongating RNA polymerase II by a mouse IgM antibody exhibits no obvious cross-talk from the detection of recruited RNA polymerase II by a rat IgG antibody. The staining protocol and image acquisition procedure were kept constant across all samples. Look-up tables for image display are identical for all immunofluorescence channels for direct comparison of intensities. Similar results were observed for *N* = 2, 3, 3 embryos per condition, viewing *n* = 18, 30, 30 different positions in these embryos.

**Fig. 11:**
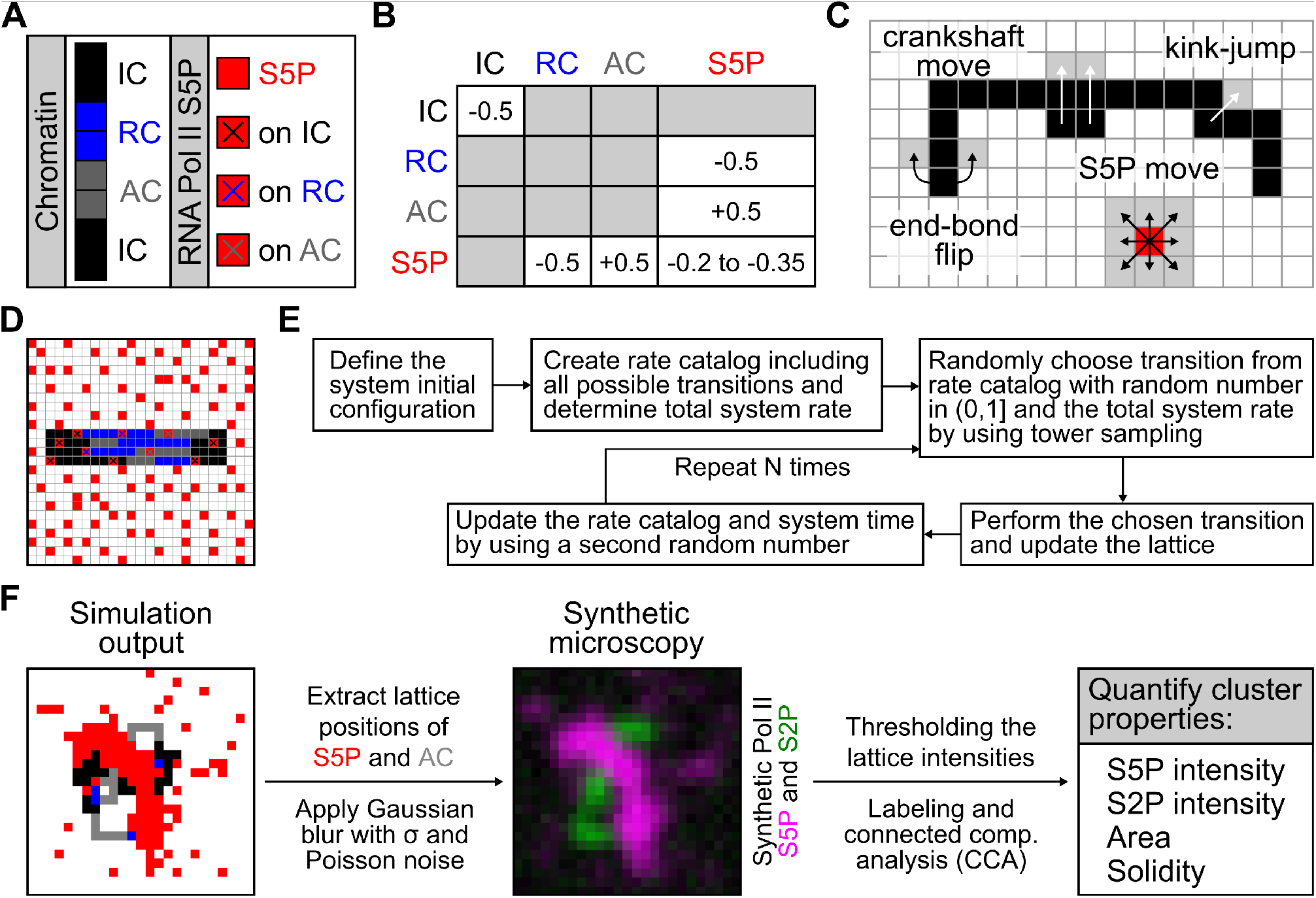
Theoretical model of surface condensation. **A)** Model components: Chromatin as polymer with different subregions and S5P as single lattice sites. **B)** Interspecies affinities: Negative values are attraction, positive ones repulsion and gray sectors are neutral. **C)** Move set: Verdier-Stockmayer move set for polymers, including end-bond flip, kink-jump and crankshaft move. Single S5P lattice sites can move to all eight nearest neighbouring sites. **D)** Lattice initial configuration: Stack of four polymers (*N_Polymer_* = 4) of length *L_Polymer_* = 20 within a 25*×*25 lattice. Single S5P (*N_S_*_5_*_P_* = 100) are randomly distributed within the lattice. **E)** General simulation framework for the lattice kinetic Monte-Carlo (LKMC) simulations. **F)** Analysis pipeline of simulation results including production of synthetic microscopy images and quantification of cluster properties.

